# Acoustic Transparency Enabling Functional Ultrasound Imaging Through Mouse and Human Skulls

**DOI:** 10.1101/2025.08.22.671878

**Authors:** Zikai Wang, Zhengchang Kou, Junhang Zhang, Jingcheng Li, Chen Gong, Chutian Lin, Jincheng Wu, Michael L Oelze, Qifa Zhou, Yaoheng Yang

**Author notes:** **Correspondence:** Yaoheng Yang.

## Abstract

Ultrasound is one of the most widely used non-invasive imaging modalities for internal organs, yet its application to the brain has been hindered for decades by the skull’s acoustic barrier. This same barrier has kept functional ultrasound imaging (fUSI)—an emerging technology capable of capturing brain-wide neural activity in real time at sub-100 μm resolution—from reaching its transformative potential in neuroscience and clinical medicine. Here, we present an acoustic transparency strategy by modulating skull acoustic properties, to render the brain visible to ultrasound. A brief topical application of an FDA-approved chelating agent matches the skull’s acoustic impedance and sound speed to those of soft tissue, enabling nearly complete ultrasound transmission (94.0 ± 4.4%) with minimal energy loss or distortion. Leveraging this discovery, we developed an acoustic-transparent fUSI platform that maps brain activation across the full brain depth at ∼20 μm spatial resolution without skull removal. This method enables safe, longitudinal brain imaging, is fully reversible, and is demonstrated to be applicable to the ex vivo human skull. This conceptually distinct paradigm—controlling acoustic wave propagation via acoustic property modulation—offers a practical and generalizable solution to one of the most persistent obstacles in transcranial ultrasound imaging, opening the door to broader clinical and research application of ultrasound neuroimaging.

## Introduction

Advances in our understanding of the brain are often driven by technological innovation, and functional ultrasound imaging (fUSI) represents one such emerging transformative technology. fUSI enables brain-wide functional imaging with exceptional resolution—without the need for contrast agents or genetic modifications—by harnessing the highly sensitive detection of neural activity–induced cerebral blood volume (CBV) changes through neurovascular coupling (*1*, *2*). fUSI offers a distinctive combination of capabilities: sub-100 μm spatial resolution, ∼100 ms temporal resolution, and high sensitivity across wide fields of view spanning several centimeters (*3*, *4*). Such performance is unmatched by most existing neuroimaging modalities such as functional MRI (fMRI) (*4*), diffuse optical tomography (DOT) (*5*, *6*), electroencephalography (EEG) (*7*), and magnetoencephalography (MEG) (*8*). By providing mesoscale imaging that bridges macroscopic and microscopic neural dynamics (*4*, *9*), fUSI offers a unique window into brain function that has been largely inaccessible to conventional technologies. Beyond its technical strengths, fUSI’s portability, relative affordability, and broad availability have further amplified its impact, enabling applications ranging from studies in freely behaving subjects to potential daily-use scenarios (*10*). Since its introduction in 2011 (*11*), fUSI has rapidly gained traction in both basic and translational neuroscience. It has been instrumental in uncovering long-range circuits underlying sleep regulation (*9*, *12*), interoception (*13*), sensory perception (*14*, *15*), pain (*16*, *17*), and cognition (*18*), and is beginning to show promise for next-generation brain–computer interfaces (BCIs) and diagnostics potential for neurological diseases (*19–28*).

Despite its unprecedented imaging capabilities, the skull remains the principal barrier to fUSI, preventing the technology from reaching its full potential (*3*, *9*, *10*, *19*, *29*). Acting as a complex, multilayered acoustic barrier, the skull can attenuate ultrasound energy by tens of decibels and distort wavefronts via reflection, refraction, scattering, absorption, and mode conversion (*10*, *30–32*), severely degrading image quality and often necessitating craniotomy for effective imaging (*3*, *9*, *19*, *29*). This long-standing challenge, recognized since the early days of transcranial ultrasound, remains a major obstacle to translating fUSI into widespread research and clinical use (*10*, *19*, *30*, *32*). Most strategies to mitigate skull-induced aberrations have emerged from the focused ultrasound therapy field, including time-reversal acoustics, adaptive beamforming, and full-wave phase correction (*31*, *33–36*). While effective in partially restoring wavefront coherence, these methods cannot recover the substantial energy loss caused by reflection and scattering, thereby limiting their suitability for fUSI (*10*). Alternatively, ultrasound localization microscopy (ULM) with intravenously injected microbubbles (*37–40*) has shown promise for brain vascular imaging by enhancing the signal-to-noise ratio through strong microbubble scattering. However, for functional neuroimaging, ULM is constrained by the short circulation time of microbubbles (3– 5 min) (*41*) and slow imaging speed (1-5 min) (*37*)— factors detrimental to detecting second-scale neurovascular dynamics (*10*). These persistent limitations underscore the need for a fundamentally different strategy—one that enables efficient, broadband ultrasound transmission through the intact skull with minimal energy loss and aberration.

In this study, we report a skull acoustic transparency strategy by directly modulating the skull’s acoustic properties to transform the brain from ultrasonic “invisible” to “visible”. Unlike conventional approaches that attempt to compensate for skull-induced wavefront aberrations, our method directly targets the fundamental biophysical cause of ultrasound attenuation and distortion: mismatches in acoustic properties between the skull and soft tissue. The mineralized bone matrix gives the skull high acoustic impedance and sound speed, creating abrupt acoustic boundaries at interfaces with soft tissue and within its heterogeneous microstructure, which in turn cause strong reflection, refraction, and scattering of ultrasound waves. Drawing inspiration from classical acoustics principles described by Lord Rayleigh (*42*), where acoustic impedance (*Z* = ^⬚^√*Kρ*) and sound speed (*c* = ^⬚^√*Kρ*) are determined by the bulk modulus *K*, we reasoned that reducing the skull’s calcium content (a key contributor to its high bulk modulus *K* (*43*)) would lower both acoustic impedance and sound speed. Among FDA-approved calcium-chelating agents, EDTA (ethylenediaminetetraacetic acid) is widely used in clinical settings, including for hypercalcemia treatment and as a demineralizing agent in ophthalmic, dermatologic, and dental procedures (*44–49*). We hypothesized that EDTA could match the skull’s acoustic properties to those of soft tissue, thereby rendering it acoustically transparent.

We demonstrate that a brief (∼20 min), topical EDTA application at clinically relevant concentrations enables near-complete ultrasound transmission through the mouse skull with almost no energy loss or wavefront distortion, effectively rendering the skull acoustically transparent. This method unlocked the ability of ultrasound to image the brain-wide vasculature with resolution down to 20 μm through skull, which was otherwise obscured by the skull. Building on this, we developed an acoustic transparent fUSI platform, demonstrating its capability to detect stimulus-evoked brain signals across the whole depth with single-trial sensitivity and spatiotemporal specificity without skull removal. Ex vivo acoustic measurements confirmed that the improved transmission results from modulating the skull’s acoustic properties to match those of brain tissue. Notably, this acoustic transparency method is safe and fully reversible. Finally, we demonstrate the feasibility of extending this acoustic transparency approach to human skulls, suggesting its potential for clinical translation.

## Results

### Acoustic Transparency Enables 3D Imaging of Brain Microvasculature through the Skull

EDTA is an FDA-approved synthetic amino acid and calcium-chelating agent, widely used in clinical and laboratory settings to form stable, water-soluble complexes for excretion (e.g., 17% solution for root canal irrigation) (*44–49*). Here, we repurposed EDTA to modulate skull acoustic properties by topically extracting calcium, thereby matching its acoustic impedance and sound speed to those of soft tissue (**Fig. 1**). We hypothesized that skull treatment with EDTA would enhance ultrasound transmission through the skull by minimizing back-reflection, scattering, and refraction, thereby overcoming key barriers to transcranial ultrasound imaging.

**Fig. 1.**
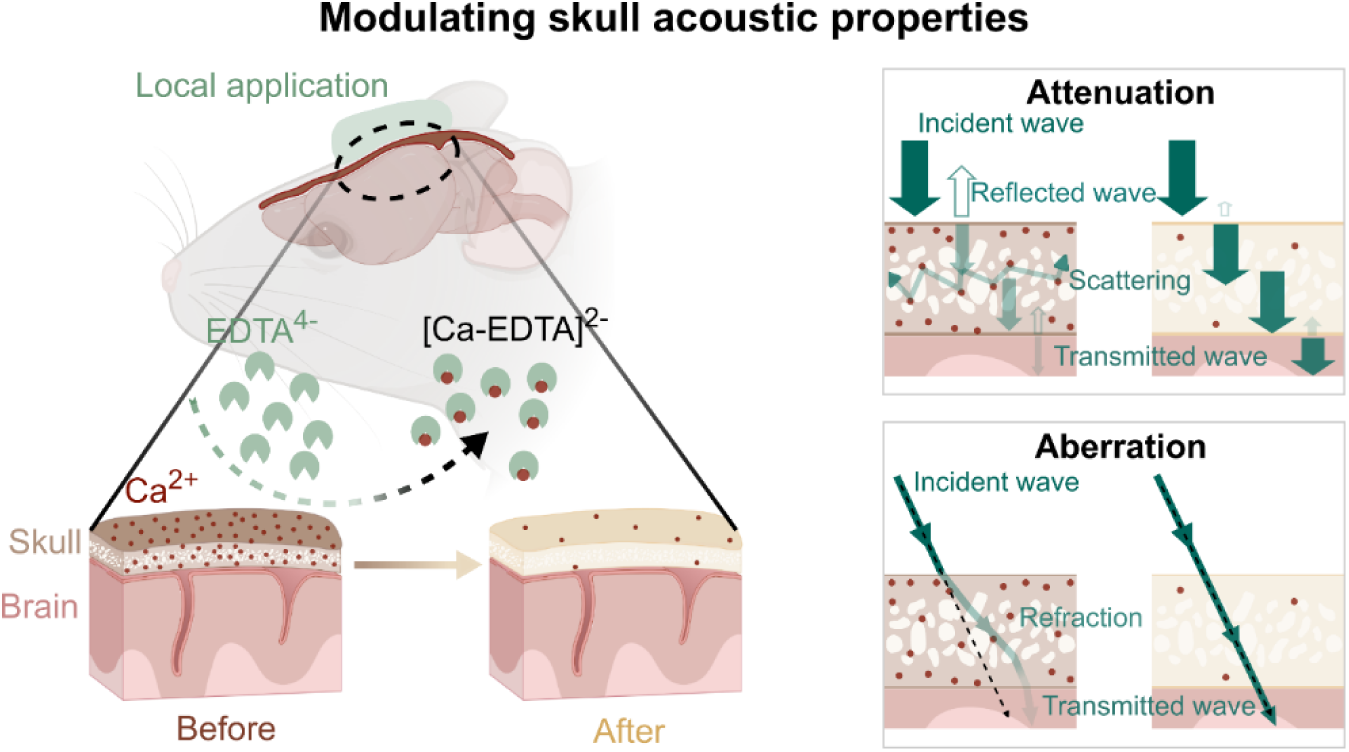
Illustration of EDTA-mediated skull acoustic transparency. EDTA chelates and extracts calcium ions (Ca²⁺) from the skull, thereby altering its acoustic properties to more closely match those of soft tissue. We hypothesize that this rematching of acoustic properties enhances ultrasound transmission and reduces wavefront aberration by minimizing back-reflection, intra-skull scattering, and refraction.

We first confirmed that EDTA, at clinically relevant concentrations (*44–46*), enables ultrasound visualization of the full-depth brain through the skull, which was otherwise nearly invisible. Using a custom-built functional ultrasound imaging (fUSI) system combining ultrafast power Doppler with null-subtraction imaging (NSI) (*11*, *50*), we performed high-resolution real-time 2D and volumetric 3D imaging of brain vasculatures without microbubbles. Before EDTA application, large portions of the brain—particularly in middle and deep regions—were almost undetectable (**Fig. 2A**). After treatment, blood vessels across the full brain depth became clearly visible (**Fig. 2B**), with a pronounced invisible-to-visible transformation in microvasculature within regions such as the thalamus and hypothalamus. This is consistent with B-mode structural imaging in which previously undetectable anatomy, such as the hippocampus, emerged after EDTA application. A positive control group with the skull removed (no-skull condition) confirmed that EDTA-treated skulls yielded images nearly indistinguishable from those obtained without a skull (**Fig. 2C**).

**Fig. 2.**
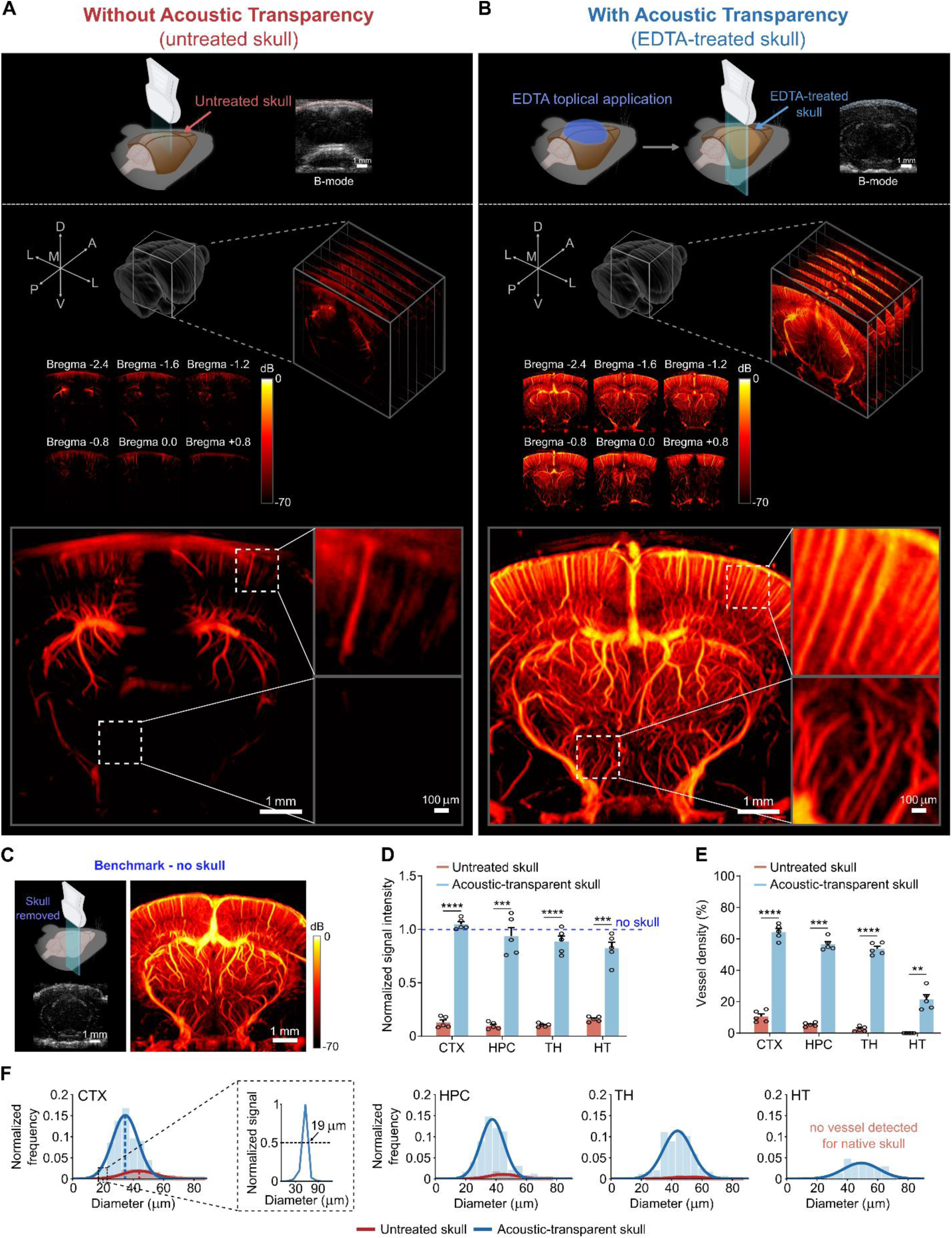
EDTA-mediated acoustic transparency enables high-resolution ultrasound imaging of brain microvasculature without skull removal. (**A-B**) Representative power Doppler images obtained before (untreated skull, A) and after EDTA treatment (acoustic-transparent skull, B) across different imaging planes. Zoomed-in views highlight microvascular structures. (**C**) Representative power Doppler images obtained after skull removal (no skull). (**D**) Quantification of power Doppler signal intensity in brains relative to the no-skull benchmark, comparing untreated skulls (n = 5) and EDTA-treated (acoustic-transparent) skulls (n = 5) across multiple brain regions (cortex, CTX; hippocampus, HPC; thalamus, TH; and hypothalamus, HT) representing different imaging depths. The blue dotted line represents the signal intensity from the no-skull group as the benchmark. (**E**) Comparison of average vessel density detected by ultrasound across the two groups: untreated skull and acoustic-transparent skull. (**F**) Histograms of the normalized frequency distribution of detected vessel diameters in different brain regions under the untreated skull and the acoustic-transparent skull, with corresponding profiles showing the minimum vessel diameter resolvable by power Doppler imaging in the cortex. Data are presented as mean ± s.e.m.; each dot represents one animal. Statistical comparisons were made using one-way ANOVA followed by Dunnett’s post hoc test. ****p < 0.0001; ***p < 0.001; **p < 0.01; *p < 0.05; NS, not significant*.

Quantitative analysis supported acoustic transparency achieved by EDTA application. We measured transcranial ultrasound signal in four regions at different depths: cortex (CTX, ∼0–1.5 mm), hippocampus (HPC, ∼1.5–2.5 mm), thalamus (TH, ∼2.5–4 mm), and hypothalamus (HT, ∼4–5.5 mm). In untreated control skulls, signals retained only 12.2 ± 1.3% of the no-skull signal intensity, with most energy lost in the bone (**Fig. 2D**). EDTA treatment enabled a trans-skull signal intensity of 94.4 ± 3.7% in average, nearly matching the no-skull condition and yielding an ∼8-fold gain over the untreated skull across all regions (8.7 ± 0.3 in CTX, 9.6 ± 0.8 in HPC, 8.6 ± 0.5 in TH, and 5.2 ± 0.4 in HT) (**Fig. 2D**). This enhancement uncovered vessels previously obscured by the skull, increasing the number of detected vessel numbers by ∼10.6-fold on average (**Fig. 2E**). Vessel density imaged after EDTA treatment was also comparable to that in the no-skull group with no significant difference. Beyond signal intensity, EDTA-mediated acoustic transparency also improved spatial resolution (**Fig. 2F**), likely by reducing skull-induced wavefront aberrations. With NSI, the smallest resolvable vessel diameter reached 19 μm in EDTA-treated mice, comparable to the no-skull group, but rarely achieved through untreated skulls. The mean vessel diameter resolved in the EDTA-treated group (36 ± 4 μm) was significantly smaller than in the untreated skull group (45 ± 5 μm) and indistinguishable from the no-skull group (37 ± 4 μm). The absence of a significant difference between the EDTA and no-skull groups confirms that EDTA treatment eliminates skull-induced degradation, achieving image quality nearly equivalent to the no-skull reference.

Collectively, these results establish proof-of-concept that topical EDTA can render the skull acoustically transparent, transforming the brain from nearly invisible to fully visible to ultrasound. This method restores signal intensity, imaging depth, and spatial resolution to near skull-free levels, enabling high-fidelity 3D visualization of brain microvasculature without skull removal.

### Acoustic-Transparent fUSI Detects Stimulus-Evoked Brain Activation Through the Skull

Beyond imaging vasculature structure, fUSI is capable of mapping neural activity by detecting associated changes in cerebral blood volume (ΔCBV) (*1*), a capability often requiring skull removal. We next tested whether EDTA-mediated acoustic transparency could enable detection of stimulus-evoked brain activation without skull removal (**Fig. 3A**). We first optimized the transparency protocol and determined that a 20-minute topical EDTA application was sufficient to achieve acoustic transparency for ultrasound transmission, while remaining brief enough to minimize potential effects on neural physiology. After treatment, we utilized fUSI-synchronized whisker stimulation to evoke brain activation. During the whisker stimulation protocol, the mouse’s right whiskers were mechanically stimulated (30 seconds per stimulation block, repeated 5 times) during the 9-minute total fUSI session. Sham controls underwent the same procedure, except the mechanical stimulator did not contact the whiskers.

**Fig. 3.**
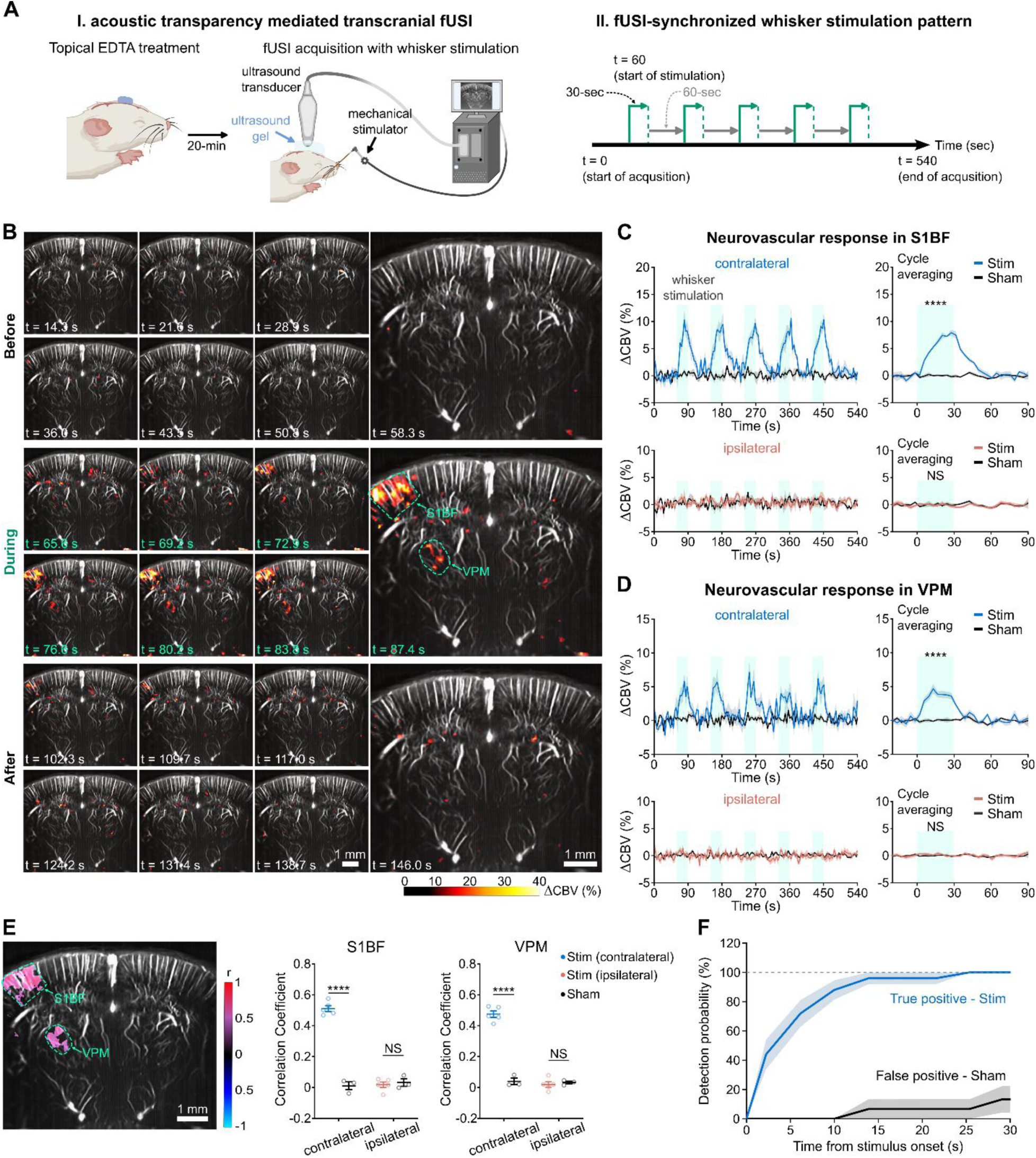
Acoustic-transparent fUSI enables reliable detection of stimulus-evoked brain activity through the skull. (**A**) Schematic of the acoustic-transparency–mediated transcranial fUSI setup for detecting whisker-stimulation–evoked brain activation. Topical EDTA treatment (∼20 min) was applied to induce skull acoustic transparency, followed by fUSI acquisition (I) synchronized with whisker stimulation (II). (**B**) Spatiotemporal maps of cerebral blood volume changes (ΔCBV) of fUSI representing neural activity in response to whisker stimulation. During the baseline period before stimulation, no significant ΔCBV changes were detected. During stimulation, ΔCBV increases were observed in the contralateral primary somatosensory barrel field (S1BF) and ventral posteromedial nucleus (VPM) of the thalamus. After stimulation, ΔCBV returned to baseline. (**C, D**) Quantification of stimulus-evoked neurovascular responses in S1BF (**C**) and VPM (**D**) comparing stimulation (n = 5) and sham (n = 3) groups. Time traces of ΔCBV across repeated stimulus cycles and cycle-averaged responses are shown for contralateral (top) and ipsilateral (bottom) brain regions. (**E**) Correlation analysis of ΔCBV signals with the stimulus, shown in a representative image (color-coded r values), along with quantified correlation coefficients in contralateral and ipsilateral S1BF and VPM for stimulation and sham groups. (**F**) Quantification of detection probability in stimulation (true positive) and sham (false positive) groups. Data are presented as mean ± s.e.m. Each dot represents one animal. Statistical comparisons were made using a two-tailed unpaired t-test; ****p < 0.0001; ***p < 0.001; **p < 0.01; *p < 0.05; NS, not significant*.

Acoustic-transparent fUSI enabled brain-wide mapping of neural activation through the skull. It reliably detected stimulation-evoked responses in the contralateral primary somatosensory cortex, barrel field (S1BF), a region known for processing whisker input (**Fig. 3B and 3C**) (*11*). These detected responses were both temporally and spatially specific: (1) ΔCBV increased from baseline to 9.2 ± 1.2% during stimulation and returned to baseline afterward, and (2) activation was primarily localized to the contralateral S1BF, which was orders of magnitude higher than the ipsilateral S1BF (ΔCBV = 0.08 ± 0.31%) (**Fig. 3B and 3C**). No significant activation was observed in sham controls (ΔCBV = 0.14 ± 0.07%). Notably, acoustic-transparent fUSI detected neural activation with high sensitivity in every single-trial stimulation. In addition to S1BF, we also observed significant activation in the contralateral ventral posteromedial nucleus of the thalamus (VPM, 5.3% ± 0.9%; **Fig. 3B and 3D**), a subcortical relay nucleus with known projections to the S1BF (*51*). No activation was detected in the ipsilateral VPM or sham conditions. To validate these findings, we generated pixel-wise Pearson correlation maps between the fUSI signal and the stimulation time course, including only pixels that showed a statistically significant correlation (*p* < 0.05) (**Fig. 3E**). Positive correlations were specifically observed in the contralateral S1BF (r = 0.51 ± 0.02) and the contralateral VPM (r = 0.48 ± 0.02), consistent with the ΔCBV spatial pattern and known somatosensory circuitry.

We further the sensitivity and specificity of acoustic-transparent fUSI for detecting stimulus-evoked brain activation by quantifying the probability of identifying S1BF responses across repeated stimulation trials (**Fig. 3F**). In the stimulation group, a neurovascular response in S1BF was reliably detected in every trial prior to the end of stimulation, resulting in a 100% detection probability (true positive), suggesting the high sensitivity and robustness of acoustic-transparent fUSI technique. In contrast, sham group showed minimal detection probability (false positive), reflecting the good specificity.

Together, these results demonstrate that acoustic-transparent fUSI enables high-sensitivity, high-specificity detection of stimulus-evoked brain activity across the full brain depth—without the need for skull removal.

### Acoustic Transparency Modulates Skull Acoustic Properties to Enable Near-Lossless Ultrasound Transmission

Our hypothesis for EDTA-mediated skull transparency is that calcium chelation alters the skull’s acoustic properties, rematching its impedance and sound speed to those of soft tissue. This rematching improves ultrasound transmission and wavefront fidelity, enabling high-resolution transcranial imaging. To test this hypothesis, we performed a series of ex vivo acoustic characterizations.

We first validated that EDTA-mediated acoustic transparency enables trans-skull ultrasound imaging in a wire phantom (**Fig. 4A**). Consistent with in vivo observations, B-mode ultrasound imaging showed that acoustic transparency treatment substantially elevated trans-skull signal intensity from 6.5 ± 1.9% (untreated skulls) to 88.5 ± 4.1% relative to the no-skull reference—an approximately 13.6-fold enhancement (**Fig. 4B**). Although not fully equivalent to the no-skull reference, the difference was not statistically significant (*p* = 0.26). Spatial resolution also improved markedly, from 33 ± 2 μm (untreated skull) to 22 ± 1 μm after acoustic transparency, closely matching the no-skull condition (21 ± 1 μm) (**Fig. 4C**). Minor discrepancies between ex vivo and in vivo measurements likely reflect differences in acoustic media (water vs brain tissue) and imaging modes (B-mode vs ultrafast power Doppler). These results confirm that acoustic transparency restores both signal intensity and resolution—two critical parameters for transcranial imaging.

**Fig. 4.**
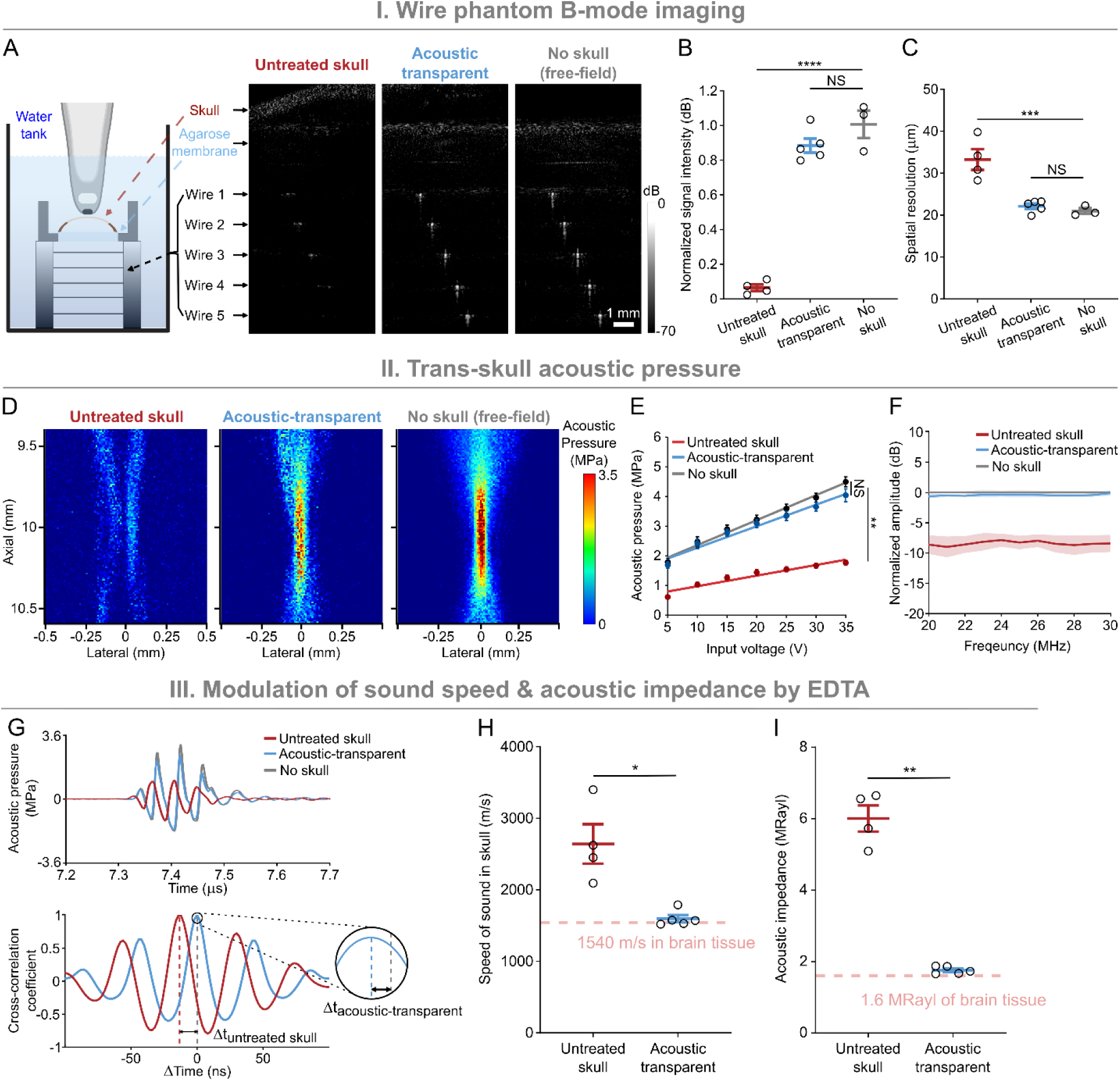
EDTA-mediated acoustic transparency re-matches skull acoustic properties to enable high-fidelity and energy lossless ultrasound transmission. (**A**) Ex vivo wire phantom experiment characterizing trans-skull ultrasound B-mode imaging under three conditions: untreated skull, acoustic-transparent skull (EDTA-treated), and no skull (free-field). B-mode images show the visibility of five embedded wires at different depths across conditions. (**B–C**) Quantification of normalized signal intensity (**B**) and spatial resolution (**C**) for each condition. (**D**) Acoustic pressure field maps captured with a capsule hydrophone show the lateral and axial distribution of transmitted ultrasound waves through untreated skull, acoustic-transparent skull, and no skull (free-field). (**E**) Measured acoustic pressure (peak positive) across a range of input voltages for the three conditions. (**F**) Frequency response curves showing the transmitted signal amplitude across a frequency range under different conditions. (**G**) Representative time-domain transmitted ultrasound waveforms collected for each condition (top), with a cross-correlation analysis to determine the time of flight relative to free field (bottom) in untreated and treated skulls. The gray dotted line indicates the reference time point for the no-skull condition (free-field). (**H–I**) Measurements of sound speed (**H**) and acoustic impedance (**I**) in untreated skulls and acoustic-transparent skulls. Pink dashed lines indicate previously reported values for brain tissue (53, 54). Data are presented as mean ± s.e.m. Each dot represents one skull from an individual mouse. p values were calculated using: (1) one-way ANOVA followed by Dunnett’s post hoc test for wire phantom and acoustic pressure characterization; and (2) a two-tailed unpaired t-test for sound speed and acoustic impedance measurements. ****p < 0.0001; ***p < 0.001; **p < 0.01; *p < 0.05; NS, not significant*.

We further demonstrated that EDTA-mediated acoustic transparency restores trans-skull ultrasound transmission and wavefront fidelity to levels approaching those of free-field propagation. Acoustic pressure fields were measured using a capsule hydrophone in a degassed water tank under three conditions—untreated skull, skull with transparency treatment, and no-skull—while keeping scanning parameters constant. Compared to the severe wavefront distortion seen with the untreated skull, acoustic transparency restored the wavefront shape to closely resemble that of the free-field (no-skull condition) (**Fig. 4D**). Acoustic transparency also increases trans-skull acoustic pressure closely approaching the free-field level (94.0 ± 4.4%) across a range of driving voltages (**Fig. 4D and 4E**). Importantly, this improvement in transmission and wavefront fidelity persisted across a broad frequency spectrum, extending up to 30 MHz (**Fig. 4F**).

By probing the skull’s acoustic properties, we found that EDTA-mediated acoustic transparency results from reductions in the skull’s sound speed and acoustic impedance, effectively rematching them to those of brain tissue. Sound speed was derived from time-of-flight (TOF) analysis, and acoustic impedance was calculated from reflection coefficients using a pulse-echo method. TOF analysis showed that ultrasound propagates significantly faster through untreated control skulls than through water, reflecting the skull’s inherently high sound speed (**Fig. 4G**). After acoustic transparency treatment, TOF increased, approaching that of water, suggesting a significant reduction in sound speed (**Fig. 4G**). Based on TOF and microCT-derived thickness, the untreated skull sound speed was 2641 ± 276 m/s (**Fig. 4H**), consistent with prior reports (*52*). EDTA treatment reduced this to 1596 ± 49 m/s—close to brain tissue (∼1540 m/s) (*53*, *54*). Acoustic impedance similarly decreased from 6.00 ± 0.37 MRayl to 1.76 ± 0.05 MRayl (**Fig. 4I**), approaching that of brain or other soft tissues (∼1.5–1.7 MRayl) (*53*, *54*). These results confirm that acoustic transparency treatment re-matches the skull’s acoustic properties to those of brain tissue, facilitating efficient ultrasound transmission.

Overall, these findings elucidate the biophysical mechanism of EDTA-mediated acoustic transparency: by reducing both acoustic impedance and sound speed, this method improves ultrasound transmission and wavefront fidelity through the skull, enabling trans-skull ultrasound brain imaging.

### EDTA-Mediated Skull Acoustic Transparency Is Safe and Reversible

As an FDA-approved compound, EDTA is considered safe when applied at controlled dosages (*44–49*). Our previous findings demonstrated that EDTA-mediated skull transparency preserved fUSI sensitivity for detecting stimulus-evoked brain activation, suggesting minimal disruption to neuronal function or neurovascular coupling. To further assess safety, we performed histological analyses of neural integrity, inflammation, and apoptosis using markers for neurons (NeuN), microglia (ionized calcium-binding adapter molecule 1, Iba1), astrocytes (glial fibrillary acidic protein, GFAP), and apoptosis (cleaved caspase-3, CC3) in brain tissues after EDTA application (**Fig. 5A**). Results revealed no significant differences in NeuN (1.6 ± 0.04 vs. 1.7 ± 0.02 × 10^3^ cells/mm^2^), Iba1 (38.8 ± 6.0 vs. 29.0 ± 3.8 cells/mm^2^), GFAP (169.3 ± 19.2 vs. 141.9 ± 10.9 cells/mm^2^), or CC3 (0.86 ± 0.24 vs. 0.71 ± 0.16 cells/mm^2^) expression between the acoustic-transparent group and untreated skull controls (**Fig. 5B**), indicating that EDTA-mediated acoustic transparency did not induce neuronal loss, neuroinflammation, or apoptosis. In contrast, the craniotomy (no-skull) group showed a significant reduction in NeuN-positive cells by 27.1 ± 4.5% and increases in Iba1 and GFAP expression by 126.2 ± 15.3% and 92.6 ± 11.5%, respectively (**Fig. 5B**), particularly in the cortex near the surgical site—consistent with acute neuroinflammatory responses and neuronal damage typically associated with surgical intervention (*55*, *56*). These findings suggest that EDTA-mediated acoustic transparency is safe for brain tissue and avoids the adverse effects associated with invasive craniotomy.

**Fig. 5.**
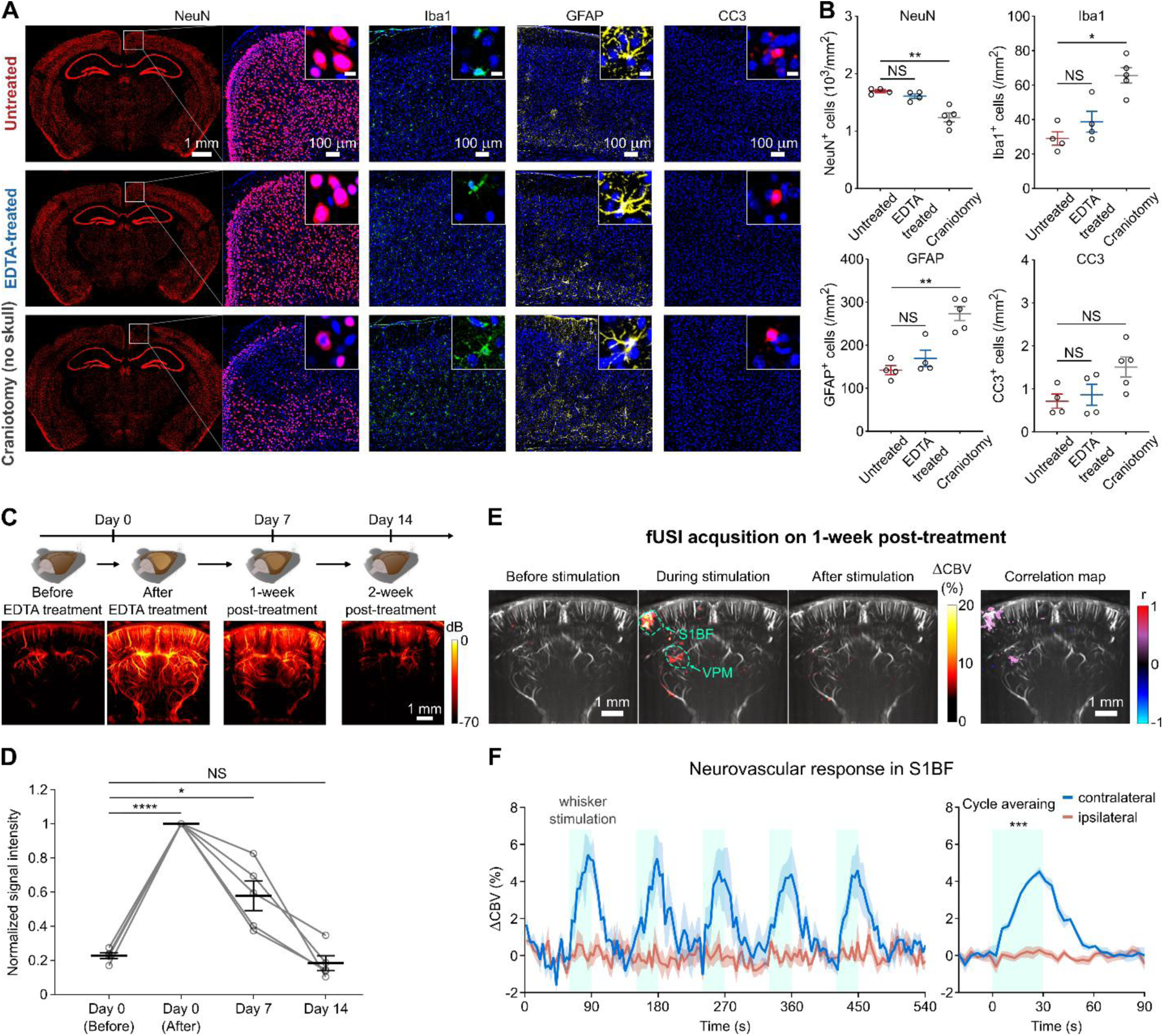
EDTA-mediated acoustic transparency is safe, reversible, and enables longitudinal fUSI. (**A**) Representative immunofluorescence images of NeuN (red), Iba1 (green), GFAP (yellow), and CC3 (magenta) under three conditions, with insets showing corresponding zoomed regions: untreated skull (n = 4), acoustic-transparent treatment (n = 4), and craniotomy (no skull, n = 5). DAPI (blue) was used for nuclear staining. (**B**) Quantification of cortical cell densities positive for each marker. (**C–D**) Longitudinal analysis of Power Doppler images (**C**) and quantification of signal intensity (**D**) at four time points (n = 5): before treatment, immediately after treatment, day 7 (1 week post-treatment), and day 14 (2 weeks post-treatment). (**E**) Representative fUSI images showing spatiotemporal maps of whisker-evoked neurovascular responses and corresponding correlation maps (right) on day 7, with both contralateral S1BF and VPM marked. (**F**) Quantification of whisker-evoked neurovascular responses in contralateral versus ipsilateral S1BF (n = 5). Time traces of ΔCBV across repeated stimulus cycles with cycle-averaged analysis are shown for both hemispheres. Data are presented as mean ± s.e.m. Each dot represents one mouse. p values were calculated using: (1) one-way ANOVA followed by Dunnett’s post hoc test for immunofluorescence quantification and longitudinal signal intensity, and (2) two-tailed paired t-test for cycle-averaged neurovascular responses in S1BF.

EDTA-mediated acoustic transparency is also reversible. Our longitudinal study found that transcranial ultrasound signal intensity peaked immediately after acoustic transparency treatment, gradually declined over one week, and returned to baseline levels comparable to pre-treatment with no significant difference (*p* = 0.93) after 2 weeks (**Fig. 5C and 5D**). This suggests that the skull recalcifies supporting the reversibility of acoustic transparency over time. Further supporting this, CT imaging—sensitive to calcium—confirmed that skull returned to its original state post-recovery. In addition, temporally and spatially specific task-evoked neurovascular responses were detected by transcranial fUSI one week after the treatment, indicating the potentiation of acoustic transparency for longitudinal investigation (**Fig. 5E and 5F**).

In summary, EDTA-mediated skull acoustic transparency is both safe and reversible, avoiding the need for invasive skull removal surgery and thereby eliminating a major barrier to the broader adoption of fUSI in basic neuroscience research and future clinical translation.

### Acoustic Transparency is Feasible in Ex Vivo Human Skull

To assess the feasibility of EDTA-mediated acoustic transparency in the ex vivo human skull, we performed experiments using extended treatment durations (120 min) and ultrasound-assisted EDTA penetration. Prior to treatment, the mouse brain imaged through the untreated ex vivo human skull was largely invisible to ultrasound (**Fig. S1A**). Following treatment, the brain vasculature became clearly visible (**Fig. S1A**). Concurrent B-mode imaging (**Fig. S1B**) confirmed the integrity of the skull, demonstrating that the improved transmission resulted from EDTA treatment rather than skull removal or acoustic wave bypass. These findings provide proof-of-concept evidence that EDTA-mediated acoustic transparency can be extended to human skulls, supporting its potential for clinical translation.

## Discussion

Overcoming the skull’s acoustic barrier has long been regarded as a longstanding grand challenge in the fields of ultrasound and neuroimaging (*10*, *19*, *29–32*). Here, we demonstrate a biophysics-inspired acoustic transparency strategy by modulating the skull’s acoustic property. Specifically, a brief topical application of EDTA reduces acoustic impedance and sound speed to values closely matching those of brain tissue, thereby enabling near-lossless transcranial ultrasound propagation. With this approach, we achieved high-resolution (∼20 μm), single-trial functional ultrasound imaging across the full brain depth through the skull. Importantly, the method is safe, reversible, and shows proof-of-concept feasibility for scaling to human skulls.

The effectiveness of EDTA-mediated acoustic transparency lies in acoustic property rematching between the skull and soft tissue. EDTA-treatment lowers the skull’s acoustic impedance from 6.00 ± 0.37 MRayl to 1.76 ± 0.05 MRayl and sound speed from 2641 ± 276 m/s to 1596 ± 49 m/s—values closely aligned with brain tissue (∼1.5–1.7 MRayl, ∼1540 m/s) (*53*, *54*). This rematching minimizes reflection, refraction at the skull–brain interface, and scattering at skull’s microstructure, thereby improving transmission and reducing wavefront aberration. As a result, ultrasound penetrates the skull with minimal energy loss or distortion, approaching free-field propagation performance.

As a conceptually distinct method, acoustic transparency directly addresses both attenuation and wavefront distortion and is well suited not only for fUSI but also for unlocking potential application across a broad range of ultrasound modalities. Conventional aberration-correction methods— such as CT-based phase correction, time-reversal acoustics, and adaptive beamforming—can partially correct wavefront distortion but are limited in recovering energy lost to reflection and scattering (*10*, *31*, *33–36*). In contrast, our method restores both acoustic transmission and wavefront integrity, approaching free-field propagation. It maintains transparency across a broad frequency range, including the high frequencies required for the spatial resolution and Doppler sensitivity essential for fUSI (*11*, *19*). Unlike contrast-enhanced ultrasound using intravenously injected microbubbles (*37–40*)—which are constrained by short circulation times and slow imaging speed—our technique enables stable, microbubble-free functional imaging with temporal resolution sufficient to capture rapid neurovascular dynamics, even at the single-trial level. By directly targeting the skull, providing broad frequency coverage, and eliminating the need for microbubbles, this approach could be extended beyond fUSI to other ultrasound modalities such as B-mode, elastography, photoacoustic and quantitative ultrasound imaging.

Transcranial fUSI, enabled by acoustic transparency, significantly expands the capabilities of the current neuroimaging toolbox and, together with existing technologies, will accelerate our understanding of brain function and improve the diagnosis of neurological disorders. Our acoustic-transparent fUSI platform allows brain-wide imaging of neural activity with sub-20 μm resolution— comparable to several neuronal diameters—combined with single-trial sensitivity and full-depth coverage through the skull. These capabilities complement existing noninvasive techniques such as fMRI, DOT, fNIRS, EEG, and MEG, each with distinct strengths. By achieving both microscopic resolution (sub-20 μm) and centimeter-scale coverage, this fUSI platform is well positioned to reveal mesoscale neural circuits and brain disease biomarkers that lie beyond the spatial resolution of large-scale functional techniques (e.g., fMRI, DOT, EEG, MEG) (*4–8*) or the field of view of implanted optical microscopy (*58*, *59*). Moreover, the ability to detect neural activity on a single-trial basis overcomes a major limitation of many hemodynamic-based imaging approaches that require averaging, enabling detection of sparse or transient neural events which are particularly valuable for brain–computer interface (BCI) development (*3*). When combined with EEG or MEG—which offer millisecond temporal resolution and direct electrical recordings (*7*, *8*)— fUSI could enhance multimodal brain-state decoding by adding deep-brain spatial information. Importantly, this approach is compatible with a wide range of ultrasound frequencies, enabling potential scalability from high-frequency imaging in small animals to lower-frequency applications in larger brains, such as nonhuman primates and potentially humans.

While promising, this proof-of-concept requires further refinement for broader application and clinical translation. First, the use of a mechanically scanned linear array for volumetric imaging limits temporal resolution for 3D imaging; integration with 2D matrix arrays (*60*, *61*) could enable real-time 3D fUSI. Second, although this study used EDTA to modulate skull acoustic properties, other FDA-approved chelating agents could, in principle, serve a similar function and warrant systematic evaluation and comparison (*62*). Finally, scaling to larger species will require optimization of EDTA dosage, application duration, and delivery strategies to accommodate thicker skulls while ensuring safety. In mice, a 20-minute topical application achieved acoustic transparency while preserving physiological responses, whereas in ex vivo human skulls, extended treatment combined with ultrasound-assisted EDTA penetration was required to enhance ultrasound transmission. Translating this approach to in vivo human applications will require substantial technological development and rigorous feasibility and safety evaluation— likely involving longer exposure times, advanced diffusion-enhancement methods, or adjuvant agents to achieve sufficient penetration without adverse effects.

In summary, we present a novel paradigm for controlling acoustic wave propagation by modulating acoustic property of the skull. By overcoming one of the most persistent barriers in transcranial ultrasound, this strategy positions fUSI for broader adoption in both neuroscience research and clinical neurology. Beyond the brain, the same principle could be applied to other organs shielded by acoustic barriers, potentially extending ultrasound’s reach in diagnostics and therapy.

## Method

### Animals

All animal procedures were approved by the Institutional Animal Care and Use Committee (IACUC) at the University of Southern California and conducted in accordance with institutional and federal guidelines. Swiss Webster mice (6–12 weeks old) were housed in a temperature-controlled (23– 26 °C) and humidity-controlled (30–70%) facility with a 12-hour light–dark cycle and *ad libitum* access to standard chow and water.

### EDTA Solution Preparation

A 20% (w/v) EDTA solution was prepared using the following procedures: First, 10 g of ethylenediaminetetraacetic acid (EDTA, E6758, Sigma-Aldrich, St. Louis, MO) was dissolved in 35 mL of Milli-Q water containing 5 mL of 10× phosphate-buffered saline (PBS, AM9625, Thermo Fisher Scientific) under magnetic stirring at room temperature. Then, ∼4 g sodium hydroxide pellets (221465, Sigma-Aldrich, St. Louis, MO) were added incrementally to facilitate dissolution. The final volume was adjusted to 50 mL with Milli-Q water, and the pH was titrated to 7.4. The solution was then sterilized by autoclaving and filtered through a 0.22 μm syringe filter (76479-024, VWR) before storage at 4 °C.

### Topical EDTA Application and Mouse Preparation

For imaging preparation, mice were initially anesthetized with 2% isoflurane for induction, followed by intraperitoneal administration of ketamine (100 mg/kg) and xylazine (20 mg/kg) for stimulus-evoked fUSI experiments, or maintained under 1–1.5% isoflurane for structural imaging. Animals were placed in a stereotaxic frame (RWD Life Science, Guangdong, China), and ophthalmic ointment (optixcare eye lube, Aventix Animal Health, Burlington, ON, CA) was applied to prevent corneal drying. Body temperature was maintained at ∼36 °C throughout the procedure. Following anesthesia, the scalp was shaved and sterilized using 70% ethanol. The scalp was incised along the midline to expose the skull, which was then cleaned using cotton swabs soaked in 3% hydrogen peroxide (216763, Sigma-Aldrich, St. Louis, MO), followed by thorough rinsing with sterile 1× PBS. The skull surface was gently abraded using autoclaved fine-grit sanding sticks to facilitate EDTA penetration. This polishing step was applied consistently across all experimental, sham, and control groups. Residual debris was removed by rinsing the surface with sterile 1× PBS.

We then applied the EDTA solution using the following protocol: The region of interest on the skull was isolated using surgical tape (Dimora paper medical tape, Winner Medical, Guangdong, China) with a central circle window to localize EDTA exposure and prevent leakage to surrounding areas. For whole-brain acoustic transparency in acute study, a commercial medical adhesive (3M Vetbond, Saint Paul, MN) was applied around the perimeter to further avoid the EDTA leakage. To deliver EDTA, sterile surgical sponges (5–10 mm diameter, customized to match the mouse skull size) were soaked in freshly prepared 20% EDTA solution and applied directly over the target area. The EDTA-soaked sponges were replaced every 5 minutes to maintain effective EDTA concentration. After the optimized treatment duration (as detailed in the Results section), the area was rinsed thoroughly with sterile 1× PBS to eliminate residual EDTA.

Subsequently, mice underwent either 3D volumetric scanning or 2D functional imaging under whisker stimulation using the established fUSI system (see following sections). Negative controls underwent identical procedures, excluding EDTA application. For positive control experiments, a cranial window (4 mm × 8 mm) was created by removing the skull over the region from −4.0 to 0.0 mm relative to bregma using a microdrill (0.6 mm drill bit, 78041, RWD Life Science, Guangdong, China).

In the longitudinal study, the skin was sutured after EDTA treatment and fUSI imaging, and the mice were allowed to recover on a warm heating pad before being returned to their home cage. 2% calcium gluconate (MWI Animal Health, Boise, ID) was locally provided to facilitate skull recalcification. Mice were re-scanned 7 days and 14 days post EDTA treatment. At the conclusion of imaging, mice were euthanized and transcranial perfused with ice-cold 1× PBS followed by 4% paraformaldehyde (PFA, Electron Microscopy Sciences, Hatfield, PA). Brains were post-fixed overnight at 4 °C in PFA for histological analysis. The skulls were harvested and stored for ex vivo acoustic characterization.

### Ultrasound Imaging System

All ultrasound imaging data were acquired using a Verasonics Vantage NXT 256 high-frequency research platform (Verasonics Inc., Kirkland, WA) equipped with a high-frequency linear array probe (L35-16vX, Vermon, France; 128 elements, 28 MHz center frequency, 0.069 mm pitch). The probe was mounted on a motorized three-axis translation stage (Velmex Inc., Bloomfield, NY) to allow precise positioning and volumetric scanning across the mouse brain.

The imaging sequences were adapted and optimized from previously established functional ultrasound protocols (*11*). For 3D vasculature structural imaging, ultrafast plane-wave imaging was performed using 19 compounded tilted plane waves ranging from −9° to +9° in 1° increments, with a pulse repetition frequency (PRF) of 15.2 kHz. This configuration yielded a compounded frame rate of 800 Hz. Power Doppler imaging (PDI) images were reconstructed by integrating 1000 compounded frames. For stimulus-evoked functional imaging, we employed a higher-speed ultrafast sequence with 9 compounded plane waves (−8° to +8°, 2° increments) at a PRF of 27 kHz, achieving a compounded frame rate of 3,000 Hz. This higher PRF and narrower angular compounding were chosen to enhance temporal resolution and better capture rapid hemodynamic responses during sensory stimulation. PDI frames were reconstructed by integrating 800 compounded frames, resulting in an effective functional frame rate of 3.6 s per frame. Note that with higher GPU computing power, the Doppler frame rate could potentially be reduced to ∼260 ms per frame.

To ensure experimental consistency and eliminate confounding acoustic variability, all acquisitions were performed with a constant probe driving voltage of 35 V across all experimental conditions.

### In Vivo Ultrasound Imaging and Whisker Stimulation

Following preparation, the ultrasound transducer was positioned approximately 3 mm above the exposed skull surface using a layer of degassed ultrasound gel (Aquasonic 100, Parker Laboratories Inc, Fairfield, NJ) to ensure acoustic coupling. For 3D vascular mapping, the probe was aligned in the coronal plane and moved along the anterior–posterior axis using a motorized linear stage, covering a total range of 7.5 mm (from −5.0 mm to +2.5 mm relative to bregma) with a 100 μm step size. Additional 3D scans were acquired in the sagittal plane by translating the probe from −4.0 mm to +4.0 mm relative to the midline with the same step size.

For stimulus-evoked fUSI, the probe was positioned in the coronal orientation and aligned to the somatosensory barrel field cortex (S1BF), approximately 1.2 mm posterior to bregma. Positioning of the probe was guided by real-time fUSI, using the hippocampal formation as a reliable anatomical landmark, compared to the standard brain atlas.

Whisker stimulation was delivered to the right whiskers using a motor-actuated rod synchronized with the ultrasound acquisition system via an external TTL trigger. The stimulation protocol consisted of the following sequence: (1) a 60-second baseline recording, (2) five stimulation blocks each comprising 30 seconds of whisker stimulation (10 Hz, 1 cm displacement) followed by 60 seconds of rest, and (3) a final 30-second post-stimulation baseline. The total acquisition time was 540 seconds per trial. For the sham control group, the stimulation protocol and probe positioning remained identical; however, the whisker pad was not physically contacted by the stimulation rod to control for motion artifacts and nonspecific hemodynamic changes.

### Generation of fUSI Imaging Using NSI

To generate functional ultrasound images from raw radiofrequency (RF) data, we implemented a two-step processing pipeline: (1) beamforming integrating the Null Subtraction Imaging (NSI) algorithm and (2) spatiotemporal clutter filtering via singular value decomposition (SVD), as described in previous studies (*63*, *64*). NSI enhances spatial resolution by synthetically creating a spatial null in the receive aperture. As described previously (*50*), three distinct apodizations were applied to the receive subapertures: a zero-mean (ZM) apodization with weights of +1 and –1 on opposite halves of the aperture, and two complementary direct current (DC1 and DC2) apodizations. The final image was constructed by subtracting the ZM envelope from the averaged DC1 and DC2 envelopes. This process results in a sharpened lateral point spread function and improved spatial resolution in power Doppler imaging.

Following NSI beamforming, SVD-based clutter filtering was applied to separate blood flow signals from tissue signals based on their distinct spatiotemporal characteristics(*63*, *64*). A fixed singular value cutoff (low 50 values and high 100 values) was used to extract the higher-rank signals corresponding to microvascular blood flow from the low-rank tissue components.

### Quantification and Brain Activation Mapping of fUSI Data

We quantitatively assessed the performance of fUSI under three experimental conditions: untreated skull, acoustic-transparent skull, and no skull (used as the benchmark). Analyses were performed in MATLAB (MathWorks). PDI signal intensity was quantified in defined brain regions of interest (ROIs) at various imaging depths, including the cortex (CTX), hippocampus (HPC), thalamus (TH), and hypothalamus (HT), based on the standard Mouse Brain Atlas (*65*). Signal intensity was computed as the relative to the intensity compared to the no skull group. For longitudinal study, signal intensity was computed as the relative to the intensity compared to that of right after EDTA treatment (day 0).

Vessel density was calculated using the following equation:

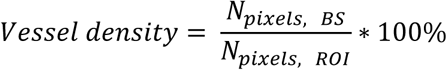

where *N_pixels, BS_* was the total number of blood signals within the ROI, and *N_pixels, BS, ROI_* was the total number of pixels in the ROI. A pixel was classified as a vessel if its intensity exceeded the local minimum plus one standard deviation within the ROI. Vessel diameter was estimated by computing the full width at half maximum (FWHM) of the signal profile across each segmented vessel, and the frequency distributions were normalized by the vessel density of each corresponding experimental condition.

Stimulus-evoked brain activation map was computed as changes in cerebral blood volume (ΔCBV) derived from power Doppler signals using custom MATLAB scripts based on established methods (*11*). For each pixel, ΔCBV was defined as the percent change in Doppler signal intensity relative to its baseline, calculated as the temporal mean over the 60 seconds prior to stimulation onset. Mean ΔCBV values were then extracted from anatomically defined ROIs, including primary somatosensory cortex, barrel field (S1BF) and ventral posteromedial nucleus of the thalamus (VPM) in both hemispheres. To generate correlation maps, we calculated the Pearson correlation coefficient between the temporal stimulation pattern and the Doppler signal time series for each pixel. Statistical significance was assessed using a two-tailed *t*-test (p < 0.05), and *p* values were corrected for multiple comparisons using the Bonferroni method for correlation analyses. Pixels meeting the corrected significance threshold were considered activated. All anatomical localizations were performed using reference to the mouse brain atlas (*65*). To quantify the probability of detecting stimulus-evoked brain activation, a detection was considered successful if the ΔCBV in the contralateral S1BF during whisker stimulation exceeded a threshold defined as the mean plus one standard deviation of the ΔCBV measured during the 30-s pre-stimulus baseline. Detection probability was then calculated cumulatively over the stimulation period and evaluated in both the stimulation and sham groups.

### Immunofluorescence Staining and Analysis

For safety analysis, mice were sacrificed 3 hours after EDTA application or after craniotomy by transcardial perfusion with ice-cold 1× PBS followed by 4% paraformaldehyde (PFA). Brains were then post-fixed overnight at 4 °C in 4% PFA. Fixed brain tissues were coronally sectioned at a thickness of 50 μm using a vibratome (VT1200, Leica Biosystems, Deer Park, IL). Sections were permeabilized in 0.3% Triton X-100 (93443, Sigma-Aldrich, St. Louis, MO) in 1× PBS (PBST) at

room temperature for 4 hours, followed by blocking in 5% normal donkey serum (NDS, 017-000-121, Jackson ImmunoResearch, West Grove, PA) in 0.1% PBST at 4 °C overnight with gentle shaking. On the following day, sections were incubated in 0.1% PBST containing primary antibodies for 6 hours at room temperature. The primary antibodies used included: chicken anti-Iba1 (1:500, Aves Labs, Davis, CA), chicken anti-GFAP (1:1000, Aves Labs, Davis, CA), rabbit anti-NeuN (1:1000, ab177487, Abcam), and rabbit anti-cleaved caspase-3 (CC3, 1:500, 9661S, Cell Signaling Technology, Danvers, MA). After three washes in 1× PBS, sections were incubated overnight at 4 °C in fluorophore-conjugated secondary antibodies (1:500, donkey anti-chicken or donkey anti-rabbit, Jackson ImmunoResearch, West Grove, PA) diluted in 1× PBS. Following additional washes, sections were mounted on glass slides with DAPI-containing antifade mounting medium (ProLong™ Diamond Antifade Mountant, Thermo Fisher Scientific) for nuclear counterstaining.

Images were acquired using a Zeiss LSM 880 inverted confocal microscope with a Plan-Apochromat 20×/0.8 NA dry objective under identical acquisition settings across all groups. Quantification of marker-positive cells was performed using QuPath software (University of Edinburgh). Positive staining was detected in the corresponding fluorescence channels, and cell counts were normalized to the region of interest (ROI) area, defined according to anatomical boundaries from the brain atlas. A consistent signal threshold was applied across all conditions to ensure comparability.

### Ex Vivo Human Skull Acoustic Transparency Experiments

We evaluated the feasibility of the acoustic transparency technique on ex vivo human temporal bone samples. A 20% EDTA solution was applied to the skull samples for 2 hours, with ultrasound treatment (40 kHz) applied concurrently to accelerate EDTA penetration, followed by a 1× PBS wash. Prior to treatment, all skull samples were stored in PBS within a degassed chamber. To assess ultrasound transmission, we performed trans-human skull power Doppler imaging using an in vivo mouse preparation covered by a human skull sample. First, acoustic transparency treatment was applied to the mouse skull as described in the previous section. The untreated or EDTA-treated human skull sample was then placed between the mouse and the ultrasound imaging probe. Power Doppler imaging was performed under three conditions: (1) without a human skull, (2) with an untreated human skull, and (3) with an EDTA-treated human skull, to visualize the mouse brain vasculature.

### Wire Phantom Characterization

A wire phantom was prepared by horizontally aligning 5 wire (20 μm diameter, 100211, Tungsten wire, California Fine Wire, Santa Barbara, CA) on a metal base. Skull samples (untreated or acoustic-transparent), harvested after in vivo imaging, were inserted between the transducer and the wire phantom and supported by a thin layer of 2% low-melt agarose (RPI, Mount Prospect, IL) to assess ultrafast plane-wave B-mode imaging. Specifically, the same linear array transducer used for in vivo imaging (L35-16vX) was mounted on a motorized 3-axis positioning system (Velmex Inc., Bloomfield, NYVelmex) and aligned perpendicularly to the 5 horizontal wires. The probe was positioned to 6 mm axially from the top wire, ensuring the imaging depth as the same in brain imaging. The ultrafast plane-wave imaging was performed using 19 compounded tilted plane waves ranging from −9° to +9° in 1° increments, with a pulse repetition frequency (PRF) of 15.2 kHz. This configuration yielded a compounded frame rate of 800 Hz for one B-mode image. The B-mode signal intensities of untreated and acoustic-transparent skulls were normalized to that of the free field water path (no skull) condition. The spatial resolution was measured by quantifying the full width at half maximum of each wire under each condition.

### Hydrophone Calibration

To quantify the acoustic effects of EDTA-mediated skull acoustic transparency, we calibrated the pressure field of the imaging transducer using a capsule hydrophone (HGL-0085, ONDA Crop. Sunnyvale, CA) in a tank filled with degassed deionized water. Skull samples (untreated or acoustic-transparent), harvested after in vivo imaging, were inserted between the transducer and hydrophone to assess ultrasound transmission efficiency, beam distortion, and sound speed. Specifically, the same linear array transducer used for imaging (L35-16vX) was mounted on a motorized 3-axis positioning system (Velmex Inc., Bloomfield, NY) and aligned perpendicularly to the hydrophone tip. The probe was programmed to emit focused pulses targeting at the hydrophone tip positioned 10 mm axially from the transducer surface. The hydrophone signals were collected at 500 MHz using a PicoScope 5000 Series oscilloscope (Pico Technology, UK). These configurations were held constant across all experimental groups. To map the 2D acoustic field, the transducer was scanned laterally and axially in 10 μm steps, covering a 1 mm × 1.2 mm field of view. The spatial distribution of peak positive pressure was computed to evaluate beam fidelity. Three conditions were tested: (1) untreated skull, (2) acoustic-transparent skull, and (3) no skull (free field control). Skull samples were placed 4 mm from the transducer to approximate the in vivo geometry. The lateral beam profile was extracted and quantified by its full width at half maximum (FWHM) to assess beam distortion due to skull interfaces. We also characterized the relationship between transducer input voltage (5–35 V) and transmitted pressure with different skull samples to measure their attenuation, and analyzed the normalized frequency spectrums of transmitted beam to quantify their attenuation relative to free field water path.

To quantify the acoustic velocity in skull samples, the sound speed (SoS) was calculated based on the difference in time-of-flight (*ΔTOF*) between transmitted ultrasound wave acquired with and without the skull sample. *ΔTOF* was measured by cross-correlation of the received signals to identify the time delay introduced by the skull compared to the water. Skull thickness (*d*) was determined from high-resolution (10 µm) micro-computed tomography (micro-CT) imaging (Phoenix Nanotom M, GE Measurement and Control Solutions, Germany). The acoustic velocity of the skull (*SoS_skull_*) was then calculated using the following equation: 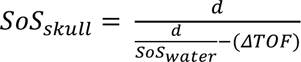 where *SoS_water_* is the known speed of sound in water (typically taken as 1480 m/s at room temperature), and *ΔTOF* is the measured time delay introduced by the skull sample relative to the water-only control. The frequency response curve was calculated by performing Fast Fourier Transform of the transmitted ultrasound signal, followed by normalizing to the no skull condition.

### Pulse-Echo Test

The acoustic impedances of untreated and acoustic-transparent skulls were calculated based on reflection coefficients measured using a pulse-echo method (*66*). The linear array imaging probe was configured in single-element mode, where each element sequentially transmitted an ultrasound pulse and received the corresponding reflected echo. Reflected signals were collected from three types of samples: untreated skull, acoustic-transparent skull, and a flat quartz plate, which served as a near-perfect reflector. All samples were aligned perpendicularly to the ultrasound probe to ensure normal incidence of the acoustic wave. The reflected pressure amplitude from the skull sample (*Vr*) was recorded and compared with the amplitude of the reflection from a quartz reference high-impedance reflector (*Vi*), which was used to approximate the incident wave amplitude. To ensure consistency, the quartz reference and skull samples positioned at the same depth.

Assuming negligible absorption and perfect reflection from the reference surface, the reflection coefficient (R) at the water–skull interface was estimated as the ratio of the reflected voltage amplitude from the skull to that from the reference reflector: 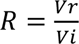. The acoustic impedance of the skull was then calculated using the standard relationship between the reflection coefficient and the acoustic impedances of two media at normal incidence:

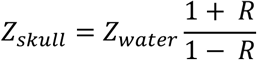

### Statistical Analysis

Statistical analyses were performed using GraphPad Prism 10 (GraphPad, San Diego, CA), except where alternative methods are indicated. Data are presented as mean ± standard error of the mean (s.e.m.) if not specified elsewhere. Comparisons between experimental groups were conducted using one-way ANOVA or two-tailed unpaired *t*-test, as appropriate. All the tests were performed after checking the normality of the data. A minimum of three independent measurements (n ≥ 3) were performed for all data, with the exact number of animals used specified in the corresponding sections. Statistical significance was defined as *p* < 0.05, with levels of significance indicated as follows: \**p* < 0.05, \*\**p* < 0.01, \*\*\**p* < 0.001, \*\*\*\**p* < 0.0001, and NS (not significant, *p* > 0.05). Data points affected by factors outside the experimental protocol, such as suboptimal cranial window quality, were excluded from the analysis to ensure accuracy and reliability of the results.

## Acknowledgments

We thank Dr. Mikhail Shapiro (Caltech) for providing the human skull and Tautis Skorka (Molecular Imaging Center, USC) for assistance with CT scanning.

## Data and Materials Availability

All raw data and code are available from the corresponding author upon reasonable request.

## Extended Data

**Fig. S1.**
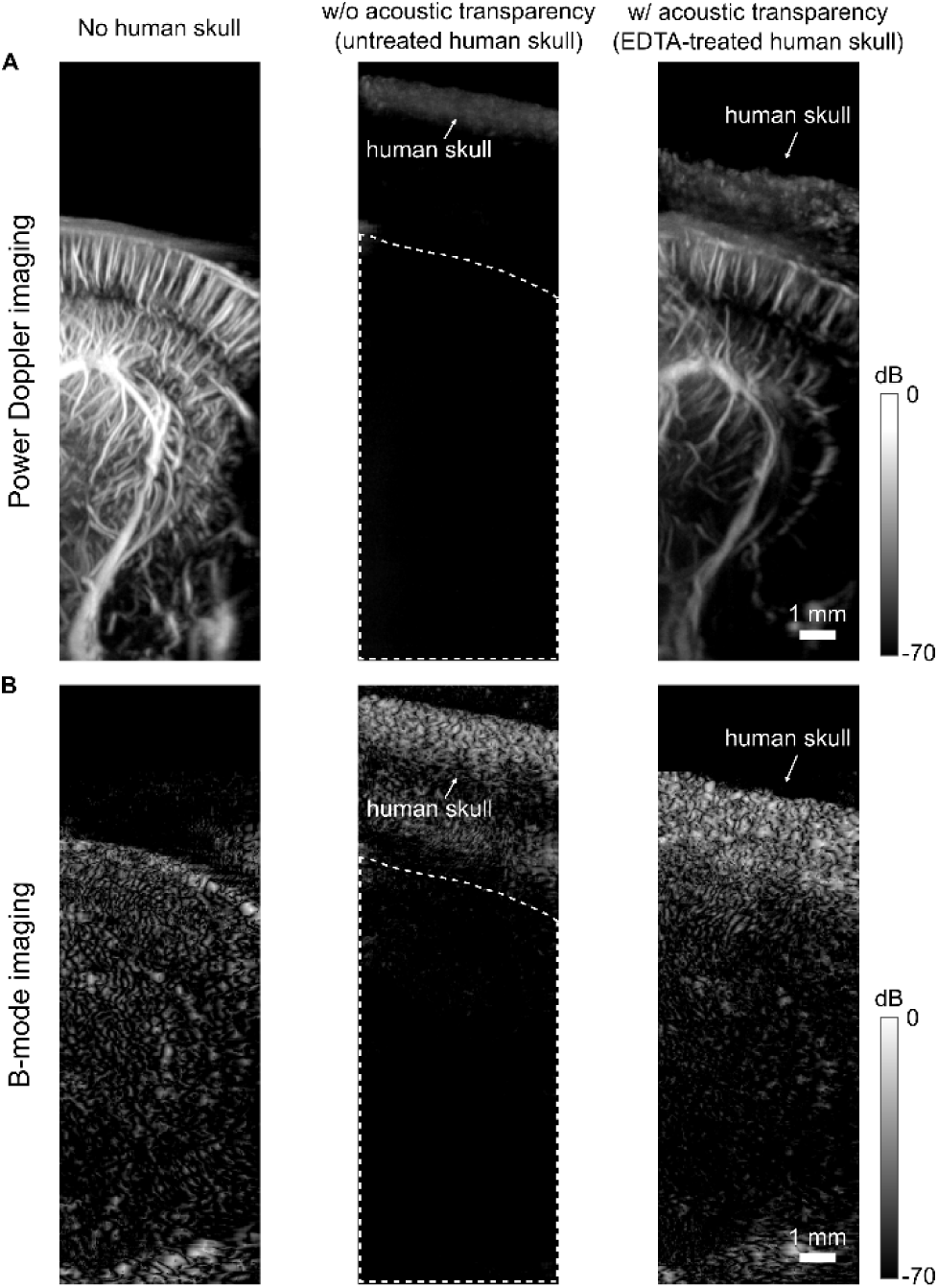
Feasibility of EDTA-mediated acoustic transparency in human skull. (**A–B**) Representative power Doppler (**A**) and B-mode (**B**) images acquired through ex vivo human skull under three conditions: (i) no human skull, (ii) untreated human skull, and (iii) EDTA-treated human skull demonstrating acoustic transparency. In all cases, the human skull was placed over an in vivo mouse brain.

## Reference

1. A.-K. Aydin, W. D. Haselden, Y. Goulam Houssen, C. Pouzat, R. L. Rungta, C. Demené, M. Tanter, P. J. Drew, S. Charpak, D. Boido, Transfer functions linking neural calcium to single voxel functional ultrasound signal. Nat Commun 11, 2954 (2020).

2. A. O. Nunez-Elizalde, M. Krumin, C. B. Reddy, G. Montaldo, A. Urban, K. D. Harris, M. Carandini, Neural correlates of blood flow measured by ultrasound. Neuron 110, 1631–1640.e4 (2022).

3. S. L. Norman, D. Maresca, V. N. Christopoulos, W. S. Griggs, C. Demene, M. Tanter, M. G. Shapiro, R. A. Andersen, Single-trial decoding of movement intentions using functional ultrasound neuroimaging. Neuron 109, 1554–1566.e4 (2021).

4. D. Boido, R. L. Rungta, B.-F. Osmanski, M. Roche, T. Tsurugizawa, D. Le Bihan, L. Ciobanu, S. Charpak, Mesoscopic and microscopic imaging of sensory responses in the same animal. Nat Commun 10, 1110 (2019).

5. A. T. Eggebrecht, S. L. Ferradal, A. Robichaux-Viehoever, M. S. Hassanpour, H. Dehghani, A. Z. Snyder, T. Hershey, J. P. Culver, Mapping distributed brain function and networks with diffuse optical tomography. Nature Photon 8, 448–454 (2014).

6. Y. Hoshi, Y. Yamada, Overview of diffuse optical tomography and its clinical applications. J. Biomed. Opt 21, 091312 (2016).

7. M. Teplan, Fundamentals of EEG Measurement. Measurement Science Review 2, 1–11 (2002).

8. M. Hämäläinen, R. Hari, R. J. Ilmoniemi, J. Knuutila, O. V. Lounasmaa, Magnetoencephalography---theory, instrumentation, and applications to noninvasive studies of the working human brain. Rev. Mod. Phys. 65, 413–497 (1993).

9. B. Li, C. Ma, Y.-A. Huang, X. Ding, D. Silverman, C. Chen, D. Darmohray, L. Lu, S. Liu, G. Montaldo, A. Urban, Y. Dan, Circuit mechanism for suppression of frontal cortical ignition during NREM sleep. Cell 186, 5739–5750.e17 (2023).

10. H. Estrada, T. Deffieux, J. Robin, M. Tanter, D. Razansky, Imaging the brain by traversing the skull with light and sound. Nat. Biomed. Eng, 1–17 (2025).

11. E. Macé, G. Montaldo, I. Cohen, M. Baulac, M. Fink, M. Tanter, Functional ultrasound imaging of the brain. Nat Methods 8, 662–664 (2011).

12. M. Matei, A. Bergel, S. Pezet, M. Tanter, Global dissociation of the posterior amygdala from the rest of the brain during REM sleep. Commun Biol 5, 1306 (2022).

13. Y. S. Zhang, D. Y. Takahashi, A. El Hady, D. A. Liao, A. A. Ghazanfar, Active neural coordination of motor behaviors with internal states. Proc. Natl. Acad. Sci. U.S.A. 119, e2201194119 (2022).

14. É. Macé, G. Montaldo, S. Trenholm, C. Cowan, A. Brignall, A. Urban, B. Roska, Whole-Brain Functional Ultrasound Imaging Reveals Brain Modules for Visuomotor Integration. Neuron 100, 1241–1251.e7 (2018).

15. H. Koorliyil, J. Sitt, I. Rivals, Y. Liu, A. Bertolo, S. Cazzanelli, A. Dizeux, T. Deffieux, M. Tanter, S. Pezet, Specific and Nonuniform Brain States during Cold Perception in Mice. The Journal of Neuroscience 44, 1 (2024).

16. L. Rahal, M. Thibaut, I. Rivals, J. Claron, Z. Lenkei, J. D. Sitt, M. Tanter, S. Pezet, Ultrafast ultrasound imaging pattern analysis reveals distinctive dynamic brain states and potent sub-network alterations in arthritic animals. Sci Rep 10, 10485 (2020).

17. T. Di Ianni, S. N. Ewbank, M. R. Levinstein, M. M. Azadian, R. C. Budinich, M. Michaelides, R. D. Airan, Sex dependence of opioid-mediated responses to subanesthetic ketamine in rats. Nat Commun 15, 893 (2024).

18. A. El Hady, D. Takahashi, R. Sun, O. Akinwale, T. Boyd-Meredith, Y. Zhang, A. S. Charles, C. D. Brody, Chronic brain functional ultrasound imaging in freely moving rodents performing cognitive tasks. Journal of Neuroscience Methods 403, 110033 (2024).

19. C. Rabut, S. L. Norman, W. S. Griggs, J. J. Russin, K. Jann, V. Christopoulos, C. Liu, R. A. Andersen, M. G. Shapiro, Functional ultrasound imaging of human brain activity through an acoustically transparent cranial window. Sci. Transl. Med. 16, eadj3143 (2024).

20. K. A. Agyeman, D. J. Lee, J. Russin, E. I. Kreydin, W. Choi, A. Abedi, Y. T. Lo, J. Cavaleri, K. Wu, V. R. Edgerton, C. Liu, V. N. Christopoulos, Functional ultrasound imaging of the human spinal cord. Neuron 112, 1710–1722.e3 (2024).

21. W. S. Griggs, S. L. Norman, T. Deffieux, F. Segura, B.-F. Osmanski, G. Chau, V. Christopoulos, C. Liu, M. Tanter, M. G. Shapiro, R. A. Andersen, Decoding motor plans using a closed-loop ultrasonic brain–machine interface. Nat Neurosci 27, 196–207 (2024).

22. C. Demene, J. Baranger, M. Bernal, C. Delanoe, S. Auvin, V. Biran, M. Alison, J. Mairesse, E. Harribaud, M. Pernot, M. Tanter, O. Baud, Functional ultrasound imaging of brain activity in human newborns. Sci. Transl. Med. 9, eaah6756 (2017).

23. A. Landemard, C. Bimbard, C. Demené, S. Shamma, S. Norman-Haignere, Y. Boubenec, Distinct higher-order representations of natural sounds in human and ferret auditory cortex. eLife 10, e65566 (2021).

24. Z. Chen, N. Li, C. Xi, J. He, J. Zhu, G. Wu, J. Xia, C. Fei, L. Sun, H. Xu, Z. Qiu, Functional Ultrasound Imaging of Auditory Responses in Comatose Patients. Research 8, 0709 (2025).

25. S. Soloukey, L. Verhoef, F. Mastik, M. Brown, G. Springeling, B. S. Generowicz, D. D. Satoer, C. M. F. Dirven, M. Smits, B. Hunyadi, S. K. E. Koekkoek, A. J. P. E. Vincent, C. I. De Zeeuw, P. Kruizinga, Mobile human brain imaging using functional ultrasound. Sci. Adv. 11, eadu9133 (2025).

26. A. Dizeux, M. Gesnik, H. Ahnine, K. Blaize, F. Arcizet, S. Picaud, J.-A. Sahel, T. Deffieux, P. Pouget, M. Tanter, Functional ultrasound imaging of the brain reveals propagation of task-related brain activity in behaving primates. Nat Commun 10, 1400 (2019).

27. J. Claron, M. Provansal, Q. Salardaine, P. Tissier, A. Dizeux, T. Deffieux, S. Picaud, M. Tanter, F. Arcizet, P. Pouget, Co-variations of cerebral blood volume and single neurons discharge during resting state and visual cognitive tasks in non-human primates. Cell Reports 42, 112369 (2023).

28. K. Blaize, F. Arcizet, M. Gesnik, H. Ahnine, U. Ferrari, T. Deffieux, P. Pouget, F. Chavane, M. Fink, J.-A. Sahel, M. Tanter, S. Picaud, Functional ultrasound imaging of deep visual cortex in awake nonhuman primates. Proc. Natl. Acad. Sci. U.S.A. 117, 14453–14463 (2020).

29. M. Tournissac, D. Boido, M. Omnès, Y. Goulam-Houssen, L. Ciobanu, S. Charpak, Cranial window for longitudinal and multimodal imaging of the whole mouse cortex. Neurophoton. 9 (2022).

30. G. Pinton, J.-F. Aubry, E. Bossy, M. Muller, M. Pernot, M. Tanter, Attenuation, scattering, and absorption of ultrasound in the skull bone. Medical Physics 39, 299–307 (2012).

31. G. T. Clement, K. Hynynen, A non-invasive method for focusing ultrasound through the human skull. Phys. Med. Biol. 47, 1219 (2002).

32. F. J. Fry, J. E. Barger, Acoustical properties of the human skull. The Journal of the Acoustical Society of America 63, 1576–1590 (1978).

33. S. Almquist, D. L. Parker, D. A. Christensen, Rapid full-wave phase aberration correction method for transcranial high-intensity focused ultrasound therapies. Journal of Therapeutic Ultrasound 4, 30 (2016).

34. B. Larrat, M. Pernot, G. Montaldo, M. Fink, M. Tanter, MR-guided adaptive focusing of ultrasound. IEEE Trans Ultrason Ferroelectr Freq Control 57, 1734–1737 (2010).

35. M. Tanter, J.-L. Thomas, M. Fink, Time reversal and the inverse filter. The Journal of the Acoustical Society of America 108, 223–234 (2000).

36. G. Kook, Y. Jo, C. Oh, X. Liang, J. Kim, S.-M. Lee, S. Kim, J.-W. Choi, H. J. Lee, Multifocal skull-compensated transcranial focused ultrasound system for neuromodulation applications based on acoustic holography. Microsyst Nanoeng 9, 45 (2023).

37. C. Errico, J. Pierre, S. Pezet, Y. Desailly, Z. Lenkei, O. Couture, M. Tanter, Ultrafast ultrasound localization microscopy for deep super-resolution vascular imaging. Nature 527, 499–502 (2015).

38. N. Renaudin, C. Demené, A. Dizeux, N. Ialy-Radio, S. Pezet, M. Tanter, Functional ultrasound localization microscopy reveals brain-wide neurovascular activity on a microscopic scale. Nat Methods 19, 1004–1012 (2022).

39. C. Demené, J. Robin, A. Dizeux, B. Heiles, M. Pernot, M. Tanter, F. Perren, Transcranial ultrafast ultrasound localization microscopy of brain vasculature in patients. Nat Biomed Eng 5, 219–228 (2021).

40. F. Bureau, L. Denis, A. Coudert, M. Fink, O. Couture, A. Aubry, Ultrasound matrix imaging for 3D transcranial in vivo localization microscopy. Sci. Adv. 11, eadt9778 (2025).

41. D. Komljenovic, T. Bäuerle, “Chapter 8 - Ultrasound Imaging of Cancer Therapy” in Cancer Theranostics, X. Chen, S. Wong, Eds. (Academic Press, Oxford, 2014), pp. 127–137.

42. J. W. S. B. Rayleigh, The Theory of Sound (Macmillan, 1896).

43. J. D. Currey, The effect of porosity and mineral content on the Young’s modulus of elasticity of compact bone. Journal of Biomechanics 21, 131–139 (1988).

44. A. Cacchio, E. De Blasis, P. Desiati, G. Spacca, V. Santilli, F. De Paulis, Effectiveness of treatment of calcific tendinitis of the shoulder by disodium EDTA. Arthritis Care & Research 61, 84–91 (2009).

45. Z. Mohammadi, S. Shalavi, H. Jafarzadeh, Ethylenediaminetetraacetic acid in endodontics. Eur J Dent 7, S135–S142 (2013).

46. I. Luque-Martinez, M. A. Muñoz, A. Mena-Serrano, V. Hass, A. Reis, A. D. Loguercio, Effect of EDTA conditioning on cervical restorations bonded with a self-etch adhesive: A randomized double-blind clinical trial. Journal of Dentistry 43, 1175–1183 (2015).

47. T. Born, C. N. Kontoghiorghe, A. Spyrou, A. Kolnagou, G. J. Kontoghiorghes, EDTA chelation reappraisal following new clinical trials and regular use in millions of patients: review of preliminary findings and risk/benefit assessment. Toxicology Mechanisms and Methods 23, 11–17 (2013).

48. T. George, M. F. Brady, “Ethylenediaminetetraacetic Acid (EDTA)” in StatPearls (StatPearls Publishing, Treasure Island (FL), 2025).

49. D. M. R. Seely, P. Wu, E. J. Mills, EDTA chelation therapy for cardiovascular disease: a systematic review. BMC Cardiovasc Disord 5, 32 (2005).

50. Z. Kou, M. R. Lowerison, Q. You, Y. Wang, P. Song, M. L. Oelze, High-Resolution Power Doppler Using Null Subtraction Imaging. IEEE Transactions on Medical Imaging 43, 3060–3071 (2024).

51. B. J. Hunnicutt, B. R. Long, D. Kusefoglu, K. J. Gertz, H. Zhong, T. Mao, A comprehensive thalamocortical projection map at the mesoscopic level. Nat Neurosci 17, 1276–1285 (2014).

52. H. Nguyen Minh, J. Du, K. Raum, Estimation of Thickness and Speed of Sound in Cortical Bone Using Multifocus Pulse-Echo Ultrasound. IEEE Trans. Ultrason., Ferroelect., Freq. Contr. 67, 568–579 (2020).

53. G. D. Ludwig, The Velocity of Sound through Tissues and the Acoustic Impedance of Tissues. The Journal of the Acoustical Society of America 22, 862–866 (1950).

54. S. Gupta, G. Haiat, C. Laporte, P. Belanger, Effect of the Acoustic Impedance Mismatch at the Bone-Soft Tissue Interface as a Function of Frequency in Transcranial Ultrasound: A Simulation and In Vitro Experimental Study. IEEE Transactions on Ultrasonics, Ferroelectrics, and Frequency Control 68, 1653–1663 (2021).

55. X. Gong, D. Mendoza-Halliday, J. T. Ting, T. Kaiser, X. Sun, A. M. Bastos, R. D. Wimmer, B. Guo, Q. Chen, Y. Zhou, M. Pruner, C. W.-H. Wu, D. Park, K. Deisseroth, B. Barak, E. S. Boyden, E. K. Miller, M. M. Halassa, Z. Fu, G. Bi, R. Desimone, G. Feng, An Ultra-Sensitive Step-Function Opsin for Minimally Invasive Optogenetic Stimulation in Mice and Macaques. Neuron 107, 38–51.e8 (2020).

56. H.-T. Xu, F. Pan, G. Yang, W.-B. Gan, Choice of cranial window type for in vivo imaging affects dendritic spine turnover in the cortex. Nat Neurosci 10, 549–551 (2007).

57. T. Selbekk, A. S. Jakola, O. Solheim, T. F. Johansen, F. Lindseth, I. Reinertsen, G. Unsgård, Ultrasound imaging in neurosurgery: approaches to minimize surgically induced image artefacts for improved resection control. Acta Neurochir 155, 973–980 (2013).

58. K. K. Ghosh, L. D. Burns, E. D. Cocker, A. Nimmerjahn, Y. Ziv, A. E. Gamal, M. J. Schnitzer, Miniaturized integration of a fluorescence microscope. Nat Methods 8, 871– 878 (2011).

59. B. A. Madruga, C. C. Dorian, L. Yang, M. Sehgal, A. J. Silva, M. Shtrahman, D. Aharoni, P. Golshani, Open-source, high performance miniature 2-photon microscopy systems for freely behaving animals. Nat Commun 16, 7125 (2025).

60. J. Sauvage, J. Poree, C. Rabut, G. Ferin, M. Flesch, B. Rosinski, A. Nguyen-Dinh, M. Tanter, M. Pernot, T. Deffieux, 4D Functional Imaging of the Rat Brain Using a Large Aperture Row-Column Array. IEEE Trans. Med. Imaging 39, 1884–1893 (2020).

61. C. Rabut, M. Correia, V. Finel, S. Pezet, M. Pernot, T. Deffieux, M. Tanter, 4D functional ultrasound imaging of whole-brain activity in rodents. Nat Methods 16, 994–997 (2019).

62. S. Entezari, S. M. Haghi, N. Norouzkhani, B. Sahebnazar, F. Vosoughian, D. Akbarzadeh, M. Islampanah, N. Naghsh, M. Abbasalizadeh, N. Deravi, Iron Chelators in Treatment of Iron Overload. Journal of Toxicology 2022, 1–18 (2022).

63. P. Song, J. D. Trzasko, A. Manduca, B. Qiang, R. Kadirvel, D. F. Kallmes, S. Chen, Accelerated Singular Value-Based Ultrasound Blood Flow Clutter Filtering With Randomized Singular Value Decomposition and Randomized Spatial Downsampling. IEEE Trans. Ultrason., Ferroelect., Freq. Contr. 64, 706–716 (2017).

64. C. Demene, T. Deffieux, M. Pernot, B.-F. Osmanski, V. Biran, J.-L. Gennisson, L.-A. Sieu, A. Bergel, S. Franqui, J.-M. Correas, I. Cohen, O. Baud, M. Tanter, Spatiotemporal Clutter Filtering of Ultrafast Ultrasound Data Highly Increases Doppler and fUltrasound Sensitivity. IEEE Trans. Med. Imaging 34, 2271–2285 (2015).

65. G. Paxinos, K. B. Franklin, The Mouse Brain in Stereotaxic Coordinates: Compact (2004).

66. D. Joshi, D. Bhatnager, A. Kumar, R. Gupta, Direct measurement of acoustic impedance in liquids by a new pulse echo technique. MAPAN 24, 215–224 (2009).

